# WAVE2 and REST/NRSF Regulate Clustered Gene Expression by Maintaining Heterochromatin Organization

**DOI:** 10.64898/2026.04.03.716287

**Authors:** Leyang Wang, Yuanxiao Tang, Haiyan Huang, Qiang Wu

**Affiliations:** Center for Comparative Biomedicine, Key Laboratory of Systems Biomedicine (MOE), Institute of Systems Biomedicine, Shanghai Jiao Tong University, Shanghai, China; Shanghai Key Laboratory of Gene Editing and Cell-based Immunotherapy for Hematological Diseases, State Key Laboratory of Medical Genomics, Ruijin Hospital, Shanghai Jiao Tong University School of Medicine, Shanghai, China

## Abstract

The actin polymerization machinery, comprising the ARP2/3 complex and its activators, the WASP family proteins, has been implicated in regulating a broad spectrum of nuclear processes, such as transcriptional regulation and nuclear organization. Here, using clustered protocadherin (*cPcdh*) and *β-globin* genes as model systems, we showed that *WAVE2*, a member of the WASP family, regulates chromatin organization by maintaining heterochromatin dynamics. Specifically, by CRISPR DNA-fragment editing, in conjunction with integrated analyses of ChIP-seq, MeDIP-seq, ATAC-seq, 4C-seq, and RNA-seq, we showed that deposition of H3K9me3, a key heterochromatin mark, is significantly decreased at the *cPcdh* locus upon *WAVE2* deletion, concurrent with aberrant accumulation of CTCF/cohesin complex at promoter regions and spatial reorganization of chromatin architecture around nucleolus. In addition, *REST/NRSF* exerts a similar heterochromatin-dependent effect on the *cPcdh* locus. Finally, genetic and genomic data showed that *WAVE2* regulates *β-globin* gene expression by maintaining heterochromatin status. Together our data suggested that *WAVE2* and *REST/NRSF* regulate clustered gene expression in a heterochromatin-dependent manner.

## Introduction

Heterochromatin, a major higher-order chromatin domain marked by histone H3 lysine 9 trimethylation (H3K9me3), is essential for nuclear organization, genome stability, and transcriptional regulation [1, 2]. At the level of 3D genome architecture, heterochromatin is segregated into distinct nuclear compartments, notably at the nuclear periphery and around the nucleolus, where it forms lamina-associated domains (LADs) and nucleolus-associated domains (NADs), respectively [3, 4]. Although both compartments are enriched in the epigenetic mark of H3K9me3 modification, they represent spatially and functionally distinct repressive environments. The Repressor Element-1 Silencing Transcription Factor (REST), also known as Neuron-Restrictive Silencer Factor (NRSF), is a master regulator of neuronal gene repression that links sequence-specific DNA binding to heterochromatin formation. Upon binding repressor elements, REST recruits corepressor complexes which lead to deposition of repressive H3K9me3 modification and removal of active chromatin marks [5]. In addition to local repression, REST contributes to higher-order genome organization by facilitating repressive nuclear compartments, including heterochromatin-rich domains and long-range promoter-enhancer interactions [6, 7].

The Arp2/3 complex and its activators, the WASP (Wiskott–Aldrich syndrome protein) family—including WAVE1, WAVE2, WAVE3, WAS, WASL, WASH, WHAMM, and JMY—are increasingly recognized as key drivers of actin polymerization [8-12]. While traditionally studied in the cytoplasm, these factors also contribute to nuclear processes [13-15]. Notably, actin polymerization has been observed at both the nuclear lamina and nucleolus both of which spatially overlap with heterochromatin domains [16-18]. However, whether members of the WASP family regulate heterochromatin organization remains largely unexplored.

Clustered genes have recently emerged as powerful model systems for revealing fundamental principles of higher-order genome organization, especially on the directionality of CTCF-mediated topological insulators and chromatin loops [19-24]. A prominent example is the human clustered protocadherin (*cPcdh*) locus, which is organized into three consecutive gene clusters—*Pcdh α, β*, and *γ* (Figure 1A) [25]. Each *cPcdh* isoform is transcribed from its own promoter located upstream of a corresponding variable exon. In the *Pcdh α* and *γ* clusters, 13 and 19 highly-similar “alternate” variable exons coexist with 2 and 3 “c-type” exons, respectively (Figure 1A) [25]. These variable exons are each *cis*-spliced to their respective set of downstream constant exons. In contrast, the *Pcdhβ* cluster comprises 16 variable exons and lacks constant regions (Figure 1A) [25]. Within the *Pcdhα* cluster, the *α1*–*α13* isoforms are expressed stochastically, whereas *αc1* and *αc2* are expressed constitutively [26, 27]. The expression of *Pcdhα* is regulated by two composite enhancers, *HS5-1* and *HS7*, which act synergistically to control transcriptional output [28-30]. Notably, the locus contains tandem convergent CTCF-binding site (CBS) elements at both promoter and enhancer regions (Figure 1A), enabling cohesin-mediated loop extrusion to establish specific promoter–enhancer contacts that underlie stochastic promoter choice [19, 31-37]. This combinatorial expression generates extensive molecular diversity of cell-surface adhesion proteins, which is essential for the assembly of complex neuronal circuits [38-45].

**FIGURE 1.**
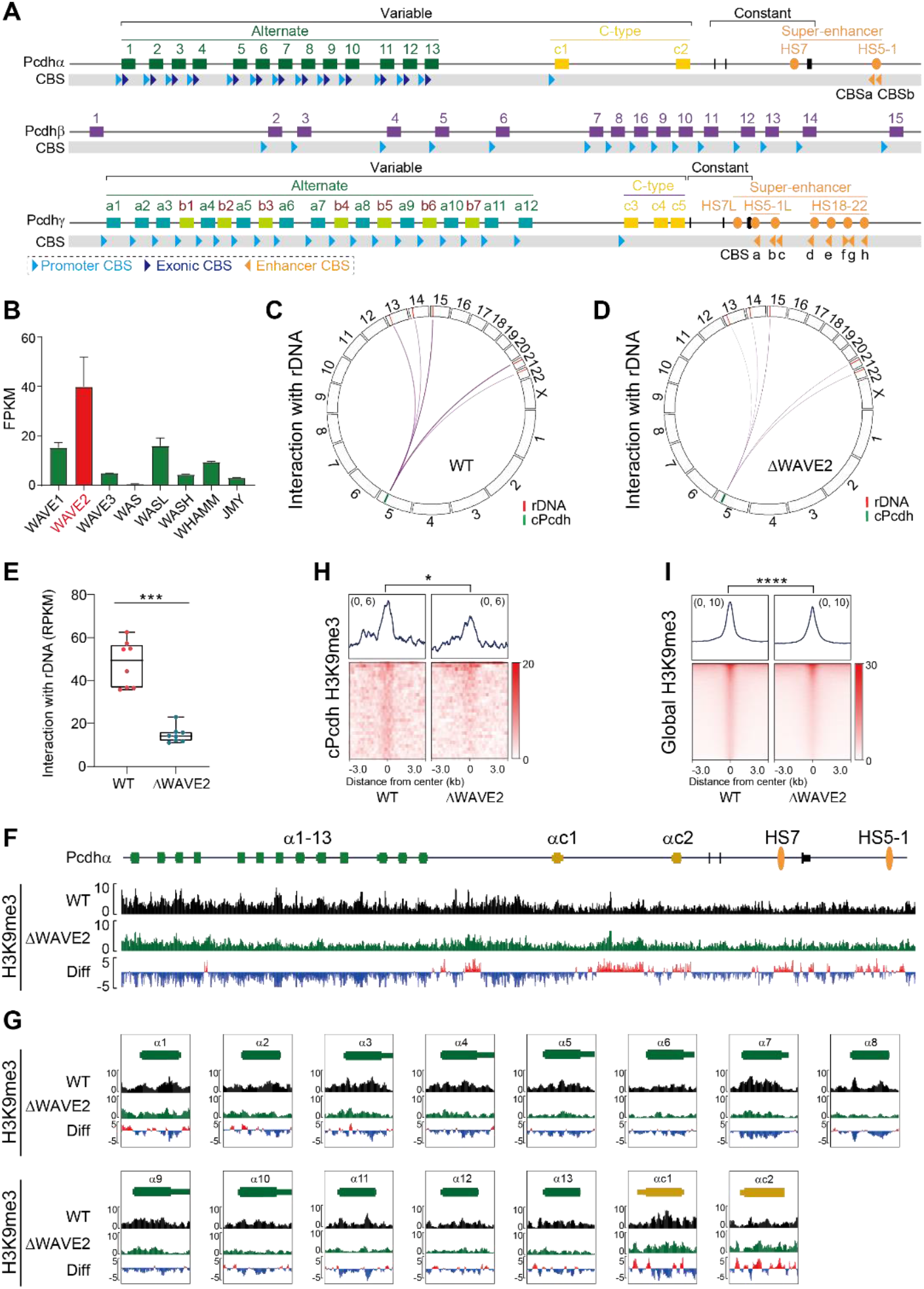
Ablation of *WAVE2* affects *cPcdh* perinucleolar organization and H3K9me3 modification. (A) Genomic architecture of the human clustered protocadherin (*cPcdh*) locus comprising three successive gene clusters. In the *Pcdhα* and *Pcdhγ* gene clusters, fifteen and twenty-two variable exons are each spliced to a single set of downstream constant exons, respectively. These variable exons are further subdivided into alternate types (*α1*-*α13*, or *γa1*-*γa12* and *γb1*-*γb7*) and ubiquitous c-types (*αc1* and *αc2, or γc3-5*). Two super-enhancers are located downstream of the *Pcdhα* and *Pcdhβγ* clusters, respectively. There are sixteen variable exons in the *Pcdhβ* cluster. The locations and orientations of CBS (CTCF Binding Site) elements are indicated below the clusters. (B) Expression levels of the members of WASP family in HEC-1-B cells (n = 2). (C and D) Circos plot representation of *cPcdh* contacts with reads mapped on rDNA in the wild-type (WT) control (C) and *WAVE2* knockout (ΔWAVE2) (D) HEC-1-B cell clones. (E) Box plot representation of *cPcdh* contacts with reads mapped on rDNA in WT and ΔWAVE2 HEC-1-B cell clones (n = 8). (F and G) ChIP-seq profiles of H3K9me3 at the *Pcdhα* gene cluster in WT and ΔWAVE2 HEC-1-B cell clones. (H and I) Aggregated peak analyses showing a significant decrease of H3K9me3 modification upon *WAVE2* deletion at the *cPcdh* locus (n = 32) (H) and genome wide (n = 27,418) (I) in HEC-1-B cells. Data are mean ± S.D, \**p* ≤ 0.05, \*\*\**p* ≤ 0.001, and \*\*\*\**p* ≤ 0.0001; unpaired two-tailed Student’s *t* test. Diff, difference (deletion vs WT); FPKM, fragments per kilobase of exon per million mapped reads; RPKM, reads per kilobase per million mapped reads.

Recent studies revealed that the *cPcdh* locus adopts a heterochromatin state in proximity to the nucleolus, with its stochastic and combinatorial expression tightly regulated by heterochromatin dynamics [35, 36, 46]. Here using CRISPR genetic deletion, in conjunction with ChIP-seq, MeDIP-seq, ATAC-seq, 4C-seq, and RNA-seq experiments, we found that WAVE2 and REST/NRSF regulate *cPcdh* gene expression via heterochromatin perinucleolar organization.

## Materials and methods

### Cell Culture and Neural Differentiation

HEC-1-B cells were cultured in MEM medium (Gibco, NY, USA) supplemented with 10% (v/v) fetal bovine serum (FBS, Sigma-Aldrich, USA), 2 mM GlutaMax (Gibco, NY, USA), 1 mM sodium pyruvate (Gibco, NY, USA), and 1% penicillin-streptomycin (Gibco, NY, USA) under standard culture conditions (37°C, 5% CO_2_, humidified atmosphere). The culture medium was replaced every day (24 h), and cells were passaged at 72 h intervals. For subculturing, adherent cells were washed with phosphate-buffered saline (PBS, Gibco, NY, USA), detached using 0.25% trypsin (Gibco, NY, USA) for 5 min at 37°C, and the enzymatic reaction was quenched with PBS containing 10% FBS. The cell suspension was centrifuged at 500 × g for 5 min (room temperature), and the cell pellet was resuspended in fresh medium for subsequent seeding.

K562 cells were cultured in DMEM (Gibco, NY, USA) supplemented with 10% FBS and 1% penicillin-streptomycin. The medium was refreshed every 24 h, and cells were passaged every 48 h. For subculturing, cells were collected without trypsin digestion and centrifuged at 500 × g for 5 min (room temperature). The pellet was resuspended in fresh medium for subsequent seeding.

Mouse E14 embryonic stem cells were cultured in MEM*α* medium (Gibco, NY, USA) supplemented with 10% (v/v) KOSR (Gibco, NY, USA), 1% FBS (Gibco, NY, USA), 10 U/ml penicillin-streptomycin (Gibco, NY, USA), 1% MEM Non-essential Amino Acid Solution (Gibco, NY, USA), 1% GlutaMax (Gibco, NY, USA), 1 mM sodium pyruvate (Gibco, NY, USA), 0.1 mM β-mercaptoethanol (Sigma-Aldrich, USA), 1 μM PD0325901 (Sigma-Aldrich, USA), 3 μM CHIR99021 (Sigma-Aldrich, USA), 10 ng/ml Lif (Sigma-Aldrich, USA). Prior to plating, cell culture plates were coated with 0.1% gelatin (Sigma-Aldrich, USA) for 20 min at room temperature. Cells were cultured at 37°C with 5% CO_2_ in a humidified atmosphere with complete medium replacement daily.

For mouse NPC differentiation [47], undifferentiated ES cells were cultured in DMEM (Gibco, NY, USA) supplemented with 15% FBS, 1% MEM Non-essential Amino Acid Solution, 2 mM L-glutamine (Gibco, NY, USA), 0.1 mM *β*-mercaptoethanol, 10 U/ml penicillin-streptomycin and 1 mM sodium pyruvate. Medium was replaced when turned yellow after pH change occurred. After 4 days, medium was replaced with the same fresh medium containing extra 5 μM retinoic acid (Sigma-Aldrich, USA). NPCs were harvested after differentiation for another 4 days.

To obtain mature neurons, NPCs were dissociated with 0.25% trypsin-EDTA and plated in DMEM/F12 (Gibco, NY, USA) supplemented with 1% N2 supplement (Gibco, NY, USA), 3 mg/mL lipid-rich bovine serum albumin (BSA) (Gibco, NY, USA), 10 U/mL penicillin-streptomycin, 3 mg/mL glucose (Gibco, NY, USA), 10 ng/mL bFGF (Gibco, NY, USA), and 1 mM L-glutamine on a tissue-cultured-treated plate. After 24 h, medium was completely replaced. On day 3, medium was replaced with a 1:1 mixture of DMEM/F12 and Neurobasal medium (Gibco, NY, USA) supplemented with 0.5% N2 supplement, 1% B27 supplement (Gibco, NY, USA), 3 mg/mL lipid-rich bovine serum albumin, 1 mM L-glutamine, and 10 U/mL penicillin-streptomycin. Medium was replaced every 2-3 days based on pH indicators. Mature neurons were harvested at day 12 of differentiation.

For human NPC differentiation, H1 ESC lines (WiCell, USA) were maintained in mTeSR Plus medium (STEMCELL, Canada) on hESC-qualified Matrigel (Corning, NY, USA)-coated plates and passaged as cell aggregates using ACCUTASE (STEMCELL, Canada). NPCs were generated from H1 cells with the STEMdiff SMADi Neural Induction Kit (STEMCELL, Canada) using the monolayer culture protocol, then maintained in the STEMdiff Neural Progenitor Medium (STEMCELL, Canada) on Matrigel (Corning, NY, USA)-coated plates under non-differentiating conditions or cryopreserved using the STEMdiff Neural Progenitor Freezing Medium (STEMCELL, Canada). NPC cells were passaged using ACCUTASE. All cells were tested negative for mycoplasma.

### CRISPR/Cas9 System for Screening Single-cell Clones

Single-guide RNA (sgRNA) expression plasmids were constructed as previously described [48]. Briefly, the pGL3-U6 vector was linearized using *Bsa* I restriction enzyme (NEB, MA, USA) to generate a cloning backbone with 5′ overhangs compatible for sgRNA insertion (TGGC and TTTG overhangs at the two ends). A pair of complementary oligonucleotides (sequences provided in Supplementary Table 1), containing the target-specific sgRNA sequence flanked by 5′ overhangs (ACCG and AAAC), were annealed and ligated into the *Bsa* I-digested vector backbone using T4 DNA ligase (NEB, MA, USA).

To screen single-cell clones, cells cultured to about 80% confluence were transfected with Lipofectamine 3000 (Invitrogen, CA, USA) in a 6-well plate with 4 μg of plasmid DNA, including 2 μg of Cas9 and 2 μg of sgRNA constructs (1 μg each one). Following 24 h incubation, WAVE2-KO cells were selected with 2 μg/ml puromycin for 4 days while REST-KO H1 cells were selected with 1 μg/ml puromycin for 4 days. Surviving cells were then serially diluted and plated in 96-well plates (for WAVE2-KO cells) or Matrigel-coated 96-plates (for REST-KO H1 cells) with approximately one cell per well to isolate single-cell clones. The plates were then cultured for a period of 14 days to let the single cell grow into a colony. The colonies were picked under a microscope and genotyped by PCR. The primer sets used are shown in Supplementary Table 1.

### Western Blot

Cultured 1 × 10^6^ cells were harvested and lysed on ice for 30 min using RIPA buffer (50 mM Tris-HCl pH 7.4, 150 mM NaCl, 1% Triton X-100, 1% sodium deoxycholate, 0.1% SDS and 1× protease inhibitors). The sample were sonicated and centrifuged at 14,000 × g for 15 min at 4°C to remove insoluble debris. Total protein concentration in the supernatant was quantified using a BCA protein assay kit (Beyotime, Shanghai, China). Protein samples were mixed with 5× SDS loading buffer containing 100 mM DTT, denatured at 95°C for 10 min, and briefly centrifuged (14,000 × g, 15 min, RT). Denatured proteins were electrophoretically resolved by SDS-PAGE on 10% polyacrylamide gels.

Resolved proteins were electrotransferred to nitrocellulose membranes. After transferring, membranes were blocked with 5% non-fat dry milk in PBST (PBS + 0.1% Tween-20) for 2 h at room temperature, followed by three times of 5 min PBST washes. Corresponding antibodies (CTCF: Millipore 07-729, MA, USA. Rad21: Abcam ab992, MA, USA. WAVE2: Sangon D262488, Shanghai, China. REST: Millipore 17-10456, MA, USA. GAPDH: Sangon D190090, Shanghai, China) were incubated with membranes overnight at 4°C with gentle agitation. After three times of 10 min PBST washes, membranes were probed with secondary antibodies (Invitrogen, CA, USA) for 90 min at room temperature. Following three times of final PBST washes (10 min each one), protein bands were visualized using an Odyssey infrared imaging system (LI-COR Biosciences, USA).

### ChIP-Seq

Cultured 5 × 10^7^ cells were harvested and crosslinked with 1% formaldehyde for 10 min at room temperature, quenched by 2 M glycine at a final concentration of 125 mM, and washed twice with ice-cold PBS. Crosslinked cells were then lysed twice in 1 ml of ChIP buffer (10 mM Tris-HCl pH 7.5, 1 mM EDTA, 1% Triton X-100, 0.1% SDS, 0.1% sodium deoxycholate, 0.15 M NaCl) supplemented with 1× protease inhibitor cocktail (Roche, BSL, Switzerland) by slowly rotating at 4°C for 10 min. Nuclei were pelleted by centrifugation (2,500 × g, 10 min, 4°C). The isolated nuclei then were resuspended in 700 μl of ChIP buffer and sonicated using a Bioruptor Sonicator (high power, 30 cycles of 30 sec ON/30 sec OFF) to shear DNA to 200–400 bp fragments. Sonicated lysates were centrifuged (14,000 × g, 10 min, 4°C), and supernatants were precleared with protein 40 μl of A/G magnetic beads (Thermo, MA, USA) for 3 h at 4°C with slow rotation. Antibody (CTCF: Millipore 07-729, MA, USA. Rad21: Abcam ab992 MA, USA. H3K9me3: Millipore 07-442, MA, USA) were added to the precleared solution and incubated overnight at 4°C with slow rotation.

Fifty μl of Protein A/G beads were added and incubated for 3 h at 4°C with slow rotation to capture antibody-protein-DNA complexes. The beads were sequentially washed with 1 ml of ChIP buffer, 1 ml of high-salt buffer (10 mM Tris-HCl pH 7.5, 1 mM EDTA, 1% Triton X-100, 0.1% SDS, 0.1% sodium deoxycholate, 0.4 M NaCl), 1 ml of no-salt buffer (high salt buffer without NaCl), 1 ml of LiCl buffer (50 mM HEPES pH 7.5, 1 mM EDTA, 1% NP-40, 0.7% sodium deoxycholate, 0.5 M LiCl) and 1 ml of 10 mM Tris-HCl (pH 7.5), by incubating at 4°C for 10 min with slow rotations, respectively. DNA-protein complexes were eluted with 400 μl of elution buffer (50 mM Tris-HCl pH 8.0, 10 mM EDTA, 1% SDS) at 65°C for 1 h with 1,000 rpm shaking. The eluted DNA-protein complex was then reverse-crosslinked overnight at 65°C with 1000 rpm shaking. Finally, the DNA-protein complex was treated with Proteinase K (NEB, MA, USA) for 2 h at 55°C, followed by RNase A (Thermo Fisher, MA, USA) for 2 h at 37°C.

DNA was purified by phenol-chloroform extraction and ethanol precipitation (2.5× volume of ethanol, 0.1× volume of 3 M NaAc (pH 5.2), and 1.5 μl of glycogen) at -80°C for 1 h. The sample was centrifuged at 14,000 × g for 10 min at 4°C. 70% ethanol was added to wash DNA pellets. The sample was centrifuged at 14,000 × g for 10 min at 4°C. After 5-min air-drying, DNA was resuspended in nuclease-free water and quantified using the Qubit fluorescent dye (Vazyme, Nanjing, China) with Qubit fluorometer (Invitrogen, CA, USA).

ChIP DNA libraries were prepared using the Universal DNA Library Prep Kit (Vazyme, Nanjing, China): DNA was first end-repaired and ligated to adapters from the Multiplex Oligos Set (Vazyme, Nanjing, China). Adapter-ligated DNA was purified with AMPure XP beads (Beckman, CA, USA) and final ChIP library was amplified by PCR. Libraries were sequenced on an Illumina platform.

### QHR-4C

Quantitative high-resolution circularized chromosome conformation capture (QHR-4C) experiments were performed as described previously with minor modifications [33]. Briefly, cultured 5 x 10^6^ cells were harvested and crosslinked with 2% formaldehyde in PBS for 10 min at room temperature with gentle rotation, followed by quenching with 200 mM glycine (final concentration) for 5 min. Crosslinked cells were washed twice with ice-cold PBS (800 × g, 5 min, 4°C) and lysed twice in 1 ml of ice-cold lysis buffer (50 mM Tris-HCl pH 7.5, 150 mM NaCl, 5 mM EDTA, 0.5% NP-40, 1% Triton X-100 and 1× protease inhibitors) for 10 min at 4°C with slow rotation. Nuclei were pelleted (800 × g, 5 min, 4°C) and resuspended in 73 μl of nuclease-free water, 10 μl of *Dpn* II buffer (NEB, MA, USA), and 2.5 μl of 10% SDS and incubated for 1 h at 37°C with shaking at 900 rpm. 12.5 μl of 20% Triton X-100 was added and incubated for 1 h at 37°C with 900 rpm shaking. Chromatin was digested with 8 μl of *Dpn* II (NEB, MA, USA) at 37°C overnight and inactivated at 65°C for 20 min with shaking at 900 rpm.

The nuclei were pelleted (1,000 × g, 1 min) and proximity ligation was performed in 100 μl of T4 ligation buffer (NEB, MA, USA) containing 2 μl of T4 DNA ligase at 16°C for 24 h. Proteinase K was added for reverse crosslinking (65°C, 4 h), followed by DNA purification via phenol-chloroform extraction and ethanol precipitation. Purified DNA was resuspended in 50 μl of nuclease-free water and sonicated with a Bioruptor Sonicator (low energy: 12 cycles of 30 sec ON/30 sec OFF) to generate 200-600 bp fragments. Finally, the concentration of the sonicated DNA was measured using the Qubit.

Linear amplification was performed with 5 μg of sonicated DNA using 5’ biotin-labeled primers for 80 cycles. Single-stranded DNA (ssDNA) was generated by heat denaturation (95°C, 5 min) and ice quenched. Then the ssDNA was captured using Streptavidin Beads (Invitrogen, CA, USA) for 2 h at room temperature. The ssDNA-bound beads were then washed twice with 200 μl of 1 x washing buffer (5 mM Tris-HCl pH 7.5, 1 M NaCl, 0.5 mM EDTA) and finally resuspended in 10 μl of nuclease-free water.

Illumina P7-compatible adapters (Supplementary Table 1) were annealed in the annealing buffer (25 mM NaCl, 10 mM Tris-HCl pH 7.5, 0.5 mM EDTA) and ligated to ssDNA at 16°C for 24 h. Libraries were amplified with two primers. The forward primer contains the Illumina P5 sequences and the sequences of the 4C viewpoint. The reverse primer contains the Illumina P7 sequences and indexes. The amplified library was sequenced on an Illumina platform. The primers for QHR-4C are listed in Supplementary Table 1.

### RNA-Seq

For each RNA-seq experiment, cultured 2 × 10^5^ cells were homogenized in 1 ml of Trizol (Invitrogen, CA, USA) and incubated for 15 min at room temperature. Following chloroform addition (0.2 ml), samples were vortexed vigorously, incubated for 3 min at room temperature, and centrifuged at 12,000 × g for 15 min at 4°C. The aqueous phase was collected and mixed with 0.5 ml of isopropanol for RNA precipitation (10 min incubation at room temperature). RNA pellets were obtained by centrifugation at 12,000 × g for 10 min at 4°C, washed with 75% ethanol, briefly air-dried, and resuspended in nuclease-free water. RNA concentration and purity (A260/A280 ratio ∼2.0) were assessed using a NanoDrop 2000 spectrophotometer. RNA-seq libraries were constructed using Universal V6 RNA-seq Library Prep Kit for Illumina (Vazyme, Nanjing, China) according to the manufacturer’s manuals.

Briefly, polyadenylated mRNA was first isolated from 1 µg of total RNA using oligo(dT)-conjugated magnetic beads (Vazyme, Nanjing, China). The purified mRNA was fragmented by heating at 94°C for 8 min. First-strand cDNA synthesis was performed by reverse transcription with random primers, followed by second-strand synthesis to generate double-stranded cDNA. After adapter ligation at both ends, the products were purified with AMPure XP beads (Beckman, CA, USA). Library amplification was carried out by PCR (98°C, 30 s for initial denaturing; 98°C, 10 s, 60°C, 30 s, 72°C, 30 s for 10 cycles; and a final extension at 72°C, 5 min) in a reaction mixture containing 2.5 μl of P5 primer, 2.5 μl of P7 primer, and 25 μl of HiFi amplification mix (Vazyme, Nanjing, China). The final libraries were sequenced on an Illumina platform.

### ATAC-Seq

For each ATAC-seq experiment, 40,000 cells were collected and washed twice with 1× PBS. Cells were then lysed in 50 μl of cold ATAC-Resuspension Buffer (RSB) (0.1% NP40, 0.1% Tween-20, and 0.01% Digitonin) by gently pipetting up and down three times. Following incubation on ice for 3 min, the lysis buffer was quenched with 1 ml of cold RSB containing 0.1% Tween-20 (without NP40 or Digitonin). Nuclei were pelleted by centrifugation at 500 × g for 10 min at 4°C. After aspirating the supernatant completely, the nuclear pellet was resuspended in 50 µl of transposition mixture via six cycles of gentle pipetting.

Chromatin fragmentation was performed by incubating the nuclei with a Tn5 transposase pre-loaded with sequencing adapters (Vazyme, Nanjing, China) at 37°C for 30 min. The reaction was terminated by adding a stopping buffer and incubating at room temperature for 5 min. DNA fragments were purified using AMPure XP beads, followed by library amplification via PCR. Finally, the libraries were sequenced on an Illumina platform.

### MeDIP-Seq

Genomic DNA was isolated from 2 × 10^5^ cells using the Wizard Genomic DNA Purification Kit (Promega, WI, USA). One μg of purified genomic DNA was fragmented to 200-600 bp using a Bioruptor Sonicator (low energy: 12 cycles of 30 sec ON/30 sec OFF) in 800 μl of TE buffer (10 mM Tris-HCl pH 7.5, 1 mM EDTA). The sheared DNA was diluted to 1 ml of immunoprecipitation (IP) buffer (15 mM Tris-HCl pH 7.5, 150 mM NaCl, 1 mM EDTA, 0.1% Triton X-100) and immunoprecipitated overnight at 4°C with 1 μg of 5-Methylcytosine (5-mC) antibody (Active Motif, 39649, CA, USA) with constant rotation. Protein A-agarose beads were added to the DNA-antibody mixture and incubated for 4 h at 4°C with rotation. The beads were washed for three times with 1× IP buffer, and methylated DNA was eluted by vigorous shaking at 55°C for 3 h in elution buffer (1% SDS, 250 mM NaCl). The immunoprecipitated DNA was purified by phenol-chloroform extraction and used for library preparation with the Universal DNA Library Prep Kit for Illumina V3 (Vazyme, Nanjing, USA) according to the manufacturer’s instructions. Finally, MeDIP-seq libraries were sequenced on an Illumina platform.

### RNA-Seq Analysis

RNA-seq raw FASTQ reads were aligned to the human reference genome (T2T-CHM13/hs1) or the mouse reference genome (mm9) using HISAT2 to generate the sequence alignment map (SAM) files. SAM files were subsequently converted into binary alignment map (BAM) files by SAMtools [49]. Transcript expression levels were then calculated in Fragments Per Kilobase of exon per Million mapped fragments (FPKM) using Cufflinks based on the BAM files [50]. Differential analysis of gene expression was performed by DEseq2 [51]. Manhattan plots illustrating the localized enrichments of upregulated genes were produced by CMplot [52]. The RNA-seq raw data of serotonergic neurons were previously published [53].

### ChIP-Seq, MeDIP-Seq, and ATAC-Seq Analysis

Raw FASTQ files were aligned to the human reference genome (T2T-CHM13/hs1) using Bowtie2 to generate SAM files. These SAM files were subsequently sorted and indexed into BAM format using SAMtools. For comparative analysis of occupancy levels across samples, the BAM files were normalized to RPKM using the bamCoverage module of deepTools with a bin size of 20 bp [54]. The resulting bedGraph files were uploaded to the UCSC Genome Browser for visualization. Narrow peaks and broad peaks were identified using MACS3 [55]. The plotHeatmap module of deepTools was used to generate heatmaps [56].

### QHR-4C Analysis

Raw FASTQ files were trimmed and PCR duplicate reads were removed by FastUniq [57]. The obtained unique reads were aligned to the human reference genome (T2T-CHM13/hs1) using Bowtie2 to generate SAM files. SAM files were converted into BAM files by SAMtools. For rDNA interaction analysis, BAM files were normalized to RPKM using the bamCoverage module of deepTools. The Circos plot was generated with the circlize package in R [58]. To analyze spatial interactions within the *Pcdhα* cluster, the BAM files were used as input for r3Cseq in the R package to calculate the reads per million (RPM) values [59]. The resulting bedGraph files were uploaded to the UCSC Genome Browser for visualization.

### Quantification and Statistical Analysis

All statistical tests were calculated using R (v4.3.3) or GraphPad (v8.4.2). Data were presented as mean ± standard deviation (SD). *p*-values were calculated using the unpaired two-tailed Student’s *t* test.

## Results

### Deletion of WAVE2 Alters *cPcdh* Perinucleolar Organization and H3K9me3 Modification

Given that actin polymerization has been observed in heterochromatin-enriched regions such as the nuclear lamina and nucleolus, we investigated whether members of the WASP family play a role in the *cPcdh* heterochromatic modification. We first performed RNA-seq experiments to assess the expression levels of members of the WASP family and found that WAVE2 is the most highly expressed member in HEC-1-B cells (Figure 1B). We then obtained a homozygous *WAVE2*-deleted cell line using CRISPR/Cas9 DNA-fragment editing through screening 116 HEC-1-B single-cell clones (Figure S1A,B) [19, 48]. The *WAVE2* knockout in the HEC-1-B cell clone was confirmed by Western blot analyses (Figure S1C).

To probe the *cPcdh* perinucleolar organization, we performed Quantitative High-Resolution Chromatin Conformation Capture Copy (QHR-4C) using *HS5-1* as a viewpoint. In wild-type cells, the *cPcdh* locus exhibits prominent spatial chromatin interactions with all regions of 45S rDNA sequences (Figure 1C), confirming the recent findings that the *cPcdh* locus is located perinucleolarly within nucleus [46]. Remarkably, the spatial chromatin interactions between *cPcdh* and nucleolus are significantly decreased upon *WAVE2* deletion (Figure 1D,E). We next performed ChIP-seq experiments using an H3K9me3-specific antibody and found that H3K9me3 depositions on almost every variable exon of the three *Pcdh* gene clusters are decreased upon *WAVE2* deletion (Figure 1F,G and S1D,E). Aggregated peak analyses showed that there is a significant decrease of H3K9me3 deposition at the *cPcdh* locus (Figure 1H). We then analyzed genome-wide H3K9me3 deposition and found that H3K9me3 depositions are also significantly decreased globally upon *WAVE2* deletion (Figure 1I). These data suggest a role of WAVE2 in *cPcdh* heterochromatin organization.

### WAVE2 Restricts CTCF/cohesin Enrichments at the *cPcdh* Locus

Since heterochromatin modification affects CTCF/cohesin enrichments [60, 61], we performed ChIP-seq experiments with a specific antibody against CTCF and found that CTCF enrichments at almost every peak within the three *Pcdh* gene clusters are increased upon *WAVE2* deletion (Figure 2A and S2A,B). Aggregated peak analyses showed that there is a significant increase of CTCF enrichments at the *cPcdh* CBS elements (Figure 2B). We then analyzed genome-wide CTCF enrichments and found that they are also significantly increased upon *WAVE2* deletion (Figure 2C).

**FIGURE 2.**
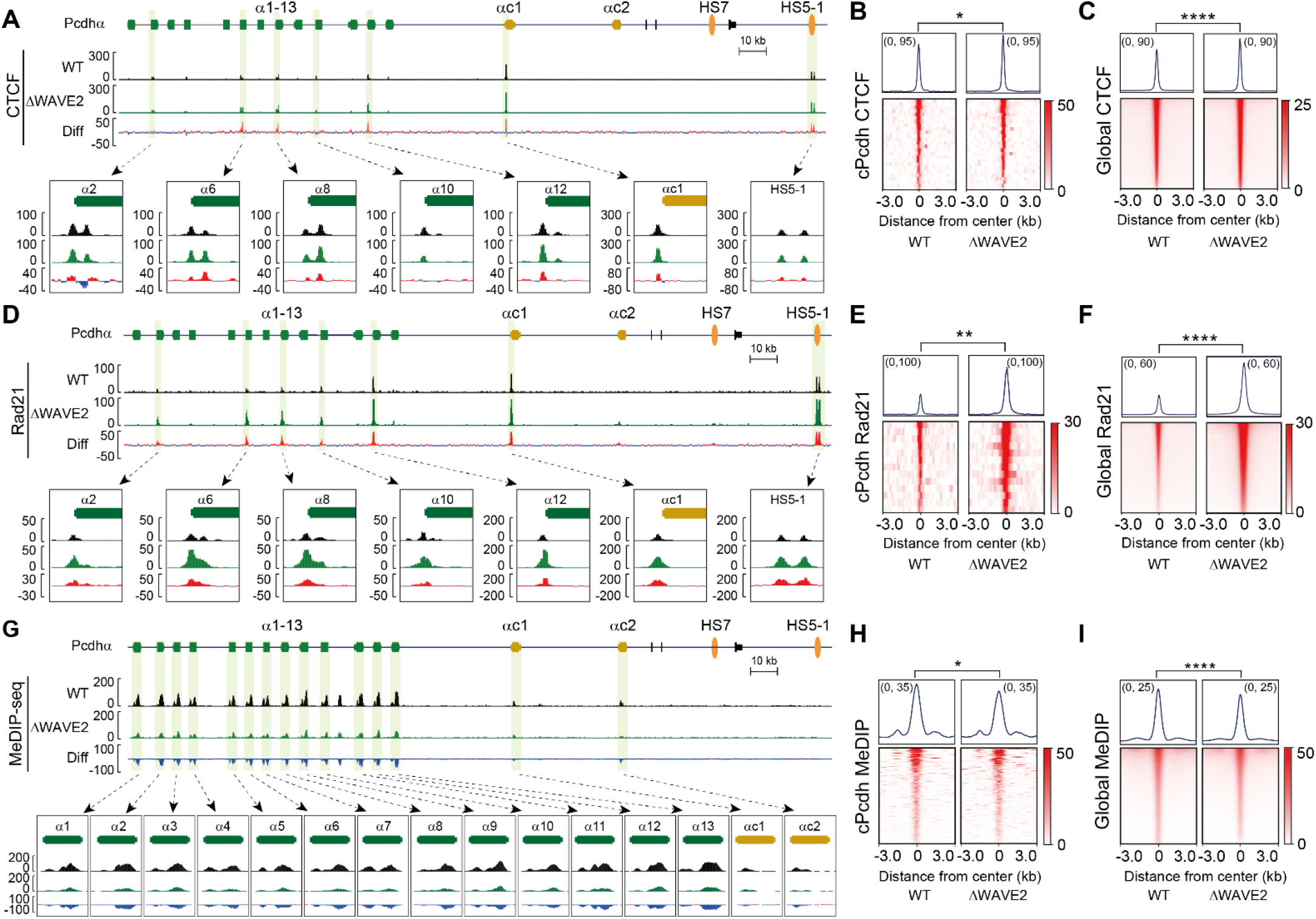
WAVE2 influences CTCF/cohesin binding profiles. (A) CTCF ChIP-seq profiles at the *Pcdhα* gene cluster in WT and ΔWAVE2 HEC-1-B cells. (B and C) Aggregated peak analyses showing a significant increase of CTCF enrichments at the *cPcdh* locus (n = 27) (B) and genome wide (n = 28,605) (C) upon *WAVE2* deletion. (D) ChIP-seq profiles indicating increased Rad21 enrichments at the *Pcdhα* gene cluster in ΔWAVE2 cells compared to WT cells. (E and F) Aggregated peak analyses showing a significant increase of Rad21 enrichments at the *cPcdh* locus (n = 14) (E) and genome-wide (n = 17,989) (F) upon *WAVE2* deletion. (G) MeDIP-seq profiles illustrating decreased DNA methylation at the *Pcdhα* gene cluster upon *WAVE2* deletion. (H and I) Aggregated peak analyses showing a significant increase of DNA methylation at the *cPcdh* locus (n = 233) (H) and genome-wide (n = 137,909) (I) upon *WAVE2* deletion. Data are mean ± S.D, \**p* ≤ 0.05, \*\**p* ≤ 0.01, and \*\*\*\**p* ≤ 0.0001; unpaired two-tailed Student’s *t* test. Diff, difference (deletion vs WT).

We next performed ChIP-seq experiments with a specific-antibody against Rad21, a subunit of the cohesin complex, and found a significant increase of Rad21 enrichments upon *WAVE2* deletion both at the *cPcdh* locus and genome wide (Figure 2D-F and S2C,D). As controls, the protein levels of CTCF and RAD21 are unchanged upon *WAVE2* deletion (Figure S2E,F). Since CTCF binding is influenced by DNA methylation, we performed Methylated DNA Immunoprecipitation Sequencing (MeDIP-seq) and observed that DNA methylation within the three *Pcdh* gene clusters is decreased upon *WAVE2* deletion (Figure 2G and S3A,B). Aggregated peak analyses showed that there is a significant decrease of DNA methylation at the *cPcdh* locus (Figure 2H). We also analyzed the genome-wide MeDIP-seq data and found that DNA methylation is significantly decreased globally upon *WAVE2* deletion (Figure 2I). Together, these data suggest that WAVE2 may restrict CTCF/cohesin enrichments by perturbing DNA methylation and H3K9me3 modification.

### WAVE2 Regulates Developmental *Pcdhα* Gene Expression

We then performed RNA-seq experiments and found an interesting switch from alternate members to ubiquitous c-type *Pcdhs* upon *WAVE2* deletion (Figure 3A). Specifically, expression levels of the two alternate members of *Pcdh α6* and *α12* are significantly decreased (Figure 3B,C), whereas expression levels of the two ubiquitous c-type of *Pcdh αc1* and *αc2* are significantly increased (Figure 3D,E). We then performed Assay for Transposase-Accessible Chromatin followed by sequencing (ATAC-seq) experiments (Figure 3F) and observed a consistent decrease of chromatin accessibility of the *α6* and *α12* promoters (Figure 3G,H) and increase of the *αc1* and *αc2* promoters (Figure 3I,J). Interestingly, the chromatin accessibility at the upstream of the *CBSa* element of the *HS5-1* enhancer is increased (Figure 3K), whereas the chromatin accessibility between the *CBSa* and *CBSb* elements is decreased (Figure 3L). To probe the chromatin interactions between distal enhancer and variable promoters, we performed QHR-4C experiments using the *HS5-1* enhancer as a viewpoint and found a consistent switch of enhancer interactions from alternate promoters to c-type *Pcdh* promoters upon *WAVE2* deletion (Figure 3M). These data suggest that *WAVE2* may regulate balanced *Pcdhα* gene expression.

**FIGURE 3.**
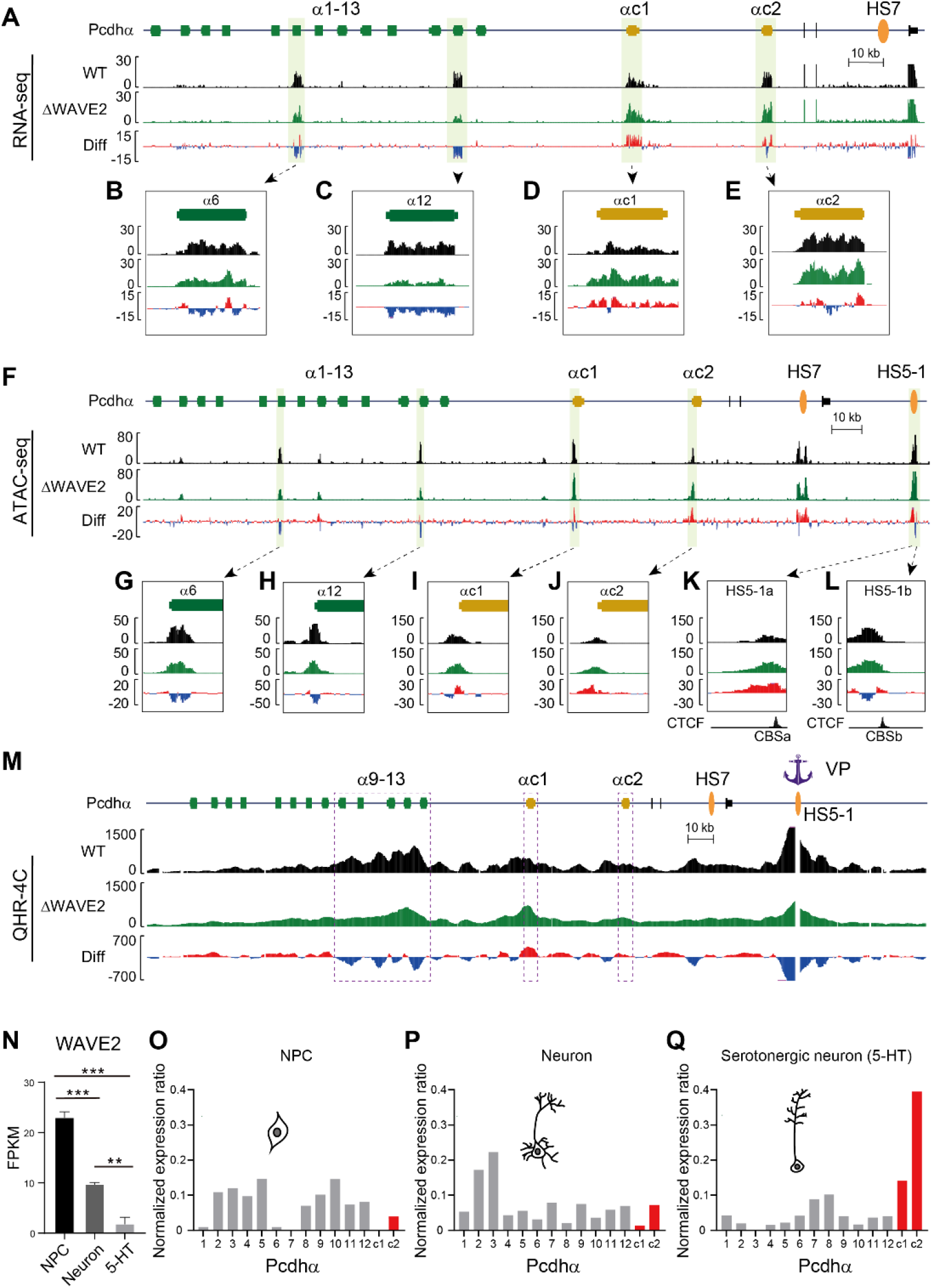
WAVE2 regulates *Pcdhα* expression. (A-E) RNA-seq profiles showing decreased expression levels of *Pcdh α6* and *α12* (B and C), but increased expression levels of *αc1* and *αc2* (D and E) upon *WAVE2* deletion. (F-L) ATAC-seq profiles showing decreased DNA accessibility at the *α6* and *α12* promoters as well as at the upstream region of the *HS5-1b* element (G, H, L) but increased DNA accessibility at the *αc1* and *αc2* promoters as well as at the upstream region of the *HS5-1a* element (I-K) in ΔWAVE2 cells compared to WT cells. (M) QHR-4C profiles using the *HS5-1* enhancer as a viewpoint in WT and ΔWAVE2 HEC-1-B cells. Differences (deletion vs WT) are shown under the 4C profiles. VP, viewpoint. (N) Expression levels of *WAVE2* in NPC (n = 3), neuron (n = 2), and serotonergic (5-hydroxytryptamine, 5-HT) neuron (n = 3). (O-Q) RNA-seq showing normalized expression ratios of *Pcdhα* isoforms in NPCs (O), neurons (P), and 5-HT (Q). Expression ratios of the ubiquitous isoforms *αc1* and *αc2* are gradually increased. Data are mean ± S.D, \*\**p* ≤ 0.01, \*\*\**p* ≤ 0.001; unpaired two-tailed Student’s *t* test. CBS, CTCF-binding site; Diff, difference (deletion vs WT); VP, viewpoint; FPKM, fragments per kilobase of exon per million mapped reads.

To further elucidate the functional role of WAVE2 in regulating *Pcdhα* isoform switch during neuronal differentiation, we established an *in vitro* differentiation system to differentiate mouse embryonic stem cells (ESCs) into neuronal progenitor cells (NPCs) and mature neurons (Figure S3C,D). We performed RNA-seq experiments to assay the developmental expression patterns of the *cPcdh* genes and observed a significant switch from alternate members to c-type genes during neural differentiation, consistent with decreased levels of *WAVE2* expression from NPCs to mature neurons (Figure 3N-Q). Together, these data suggest that WAVE2 regulates developmental *Pcdhα* gene expression.

### REST Exerts a Similar Repressive Effect on the *Pcdhα* Cluster

We previously found that REST plays a role in *Pcdhα* gene regulation [7]. To investigate the mechanism of REST in *Pcdhα* repression, we obtained a homozygous *REST*-deleted cell line using CRISPR/Cas9 DNA-fragment editing by screening 81 single-cell human ESC clones (Figure S4A-C). We then differentiated REST-depleted ESCs into NPCs (Figure S4D) and performed H3K9me3 ChIP-seq experiments to investigate heterochromatin modifications of the *cPcdh* locus. We found that H3K9me3 depositions at almost every member of the three *Pcdh* gene clusters are decreased upon *REST* deletion (Figure 4A,B and S4E,F). Aggregated peak analyses showed that there is a significant decrease of H3K9me3 deposition at the *cPcdh* locus (Figure 4C). We then analyzed genome-wide H3K9me3 modification and found that H3K9me3 deposition is significantly decreased upon *REST* deletion (Figure 4D).

**FIGURE 4.**
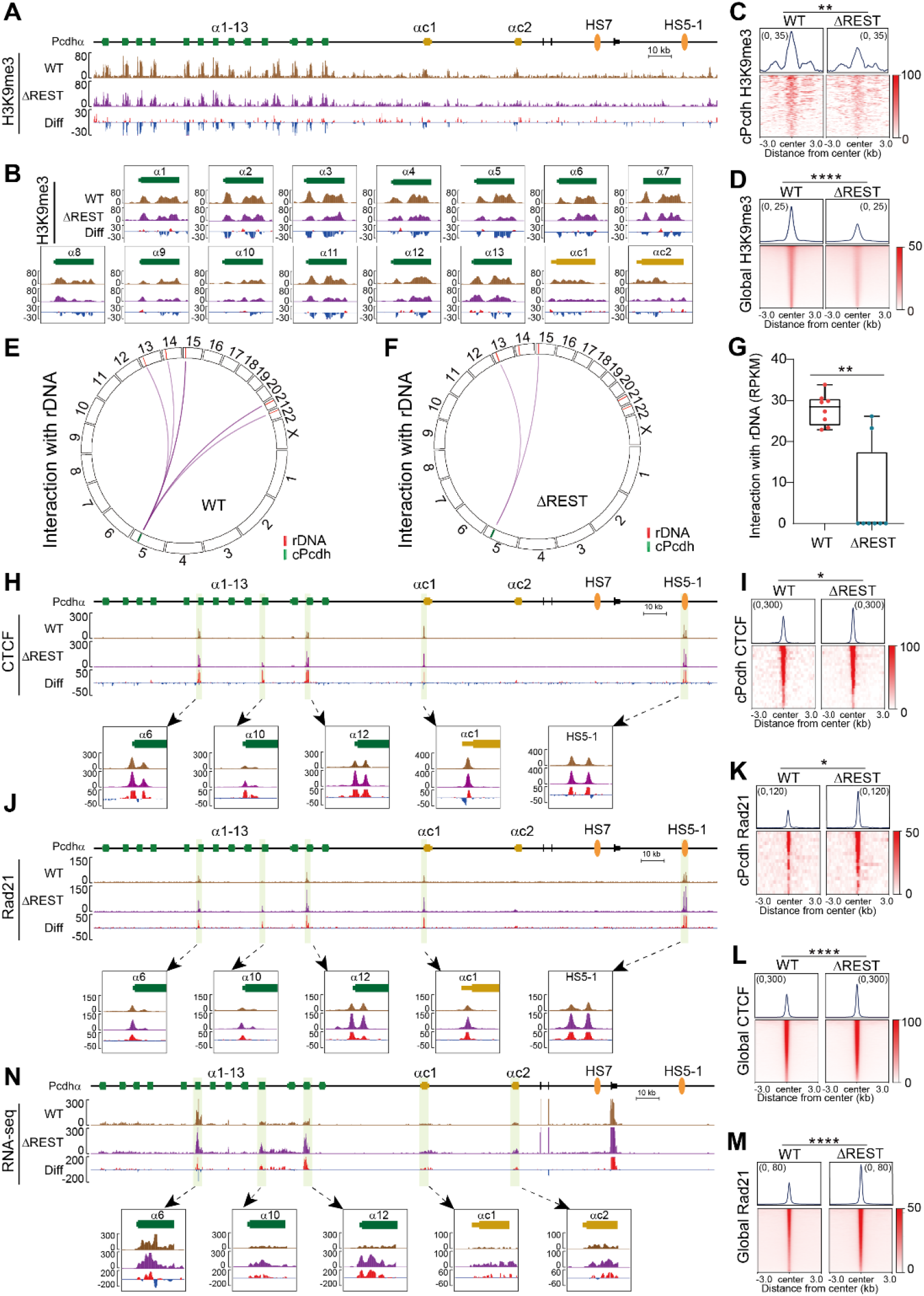
Deletion of *REST* affects *Pcdhα* perinucleolar localization and H3K9me3 modification. (A and B) ChIP-seq profiles of H3K9me3 modification at the *Pcdhα* gene cluster in NPCs. ΔREST, the *REST* knockout clone, WT, the control clone. (C and D) Aggregated peak analyses showing a significant decrease of H3K9me3 deposition at the *cPcdh* locus (n = 112) (C) and genome-wide (n = 113,514) (D) upon *REST* deletion. (E and F) Circos plot representation of *cPcdh* contacts with reads mapped on rDNA in WT (E) and ΔREST (F) cells. (G) Box plot representation of *cPcdh* contacts with reads mapped on rDNA in WT and ΔREST cells (n = 8). (H) ChIP-seq profiles showing increased CTCF enrichments at the *Pcdhα* gene cluster in ΔREST cells compared to WT cells. (I) Aggregated peak analyses showing a significant increase of CTCF enrichments at the *cPcdh* locus upon *REST* deletion (n = 22). (J) ChIP-seq profiles showing increased Rad21 enrichments at the *Pcdhα* gene cluster in ΔREST cells compared to WT cells. (K) Aggregated peak analyses showing a significant increase of Rad21 enrichments at the *cPcdh* locus upon *REST* deletion (n = 19). (L and M) Aggregated peak analyses showing a significant increase of genome-wide CTCF (n = 37,762) (L) and Rad21 (n = 28,810) (M) enrichments upon *REST* deletion. (N) RNA-seq profiles showing increased expression levels of *α6, α10, α12, αc1* and *αc2* upon *REST* deletion. Data are mean ± S.D, \**p* ≤ 0.05, \*\**p* ≤ 0.01, and \*\*\*\**p* ≤ 0.0001; unpaired two-tailed Student’s *t* test. Diff, difference (deletion vs WT); RPKM, reads per kilobase per million mapped reads.

To determine whether REST knockout affects the perinucleolar organization of the *cPcdh* locus, we performed QHR-4C using *HS5-1* as a viewpoint. In wild-type cells, the *cPcdh* locus exhibits prominent chromatin interactions with all regions of 45S rDNA sequences (Figure 4E). Remarkably, the chromatin interactions between *cPcdh* and the nucleolus were significantly decreased upon *REST* deletion (Figure 4F,G).

We next performed CTCF and Rad21 ChIP-seq experiments and found a significant increase of CTCF and Rad21 enrichments at the c*Pcdh* locus upon *REST* deletion (Figure 4H-K and S4G,H). We then analyzed genome-wide CTCF and cohesin enrichments and found that both CTCF and cohesin enrichments are significantly increased upon *REST* deletion (Figure 4L,M). Finally, we performed RNA-seq experiments and observed a significant increase of *Pcdhα* expression levels upon *REST* deletion (Figure 4N). These data suggest that *REST* regulates perinucleolar organization and represses expression of the *cPcdh* genes.

### WAVE2 Regulates Clustered *β-globin* Genes

To investigate whether WAVE2 also regulates perinuclear genes, we used the clustered *β-globin* genes as a model [62]. We note that WAVE2 is the most highly expressed member of the WASP family in K562 cells (Figure 5A). We then obtained a homozygous *WAVE2*-deleted cell line using CRISPR/Cas9 DNA-fragment editing through screening 149 single-cell K562 clones (Figure S5A,B) and carried out RNA-seq experiments in both WT and *WAVE2*-deletion single-cell clones. Interestingly, there is a localized enrichment of upregulated genes at the *β-globin* locus, with a 149-fold enrichment for upregulated transcripts compared to the rest of the genome (*P* = 3.04 × 10^-267^ by Poisson test; a 50-Kb sliding window (50Kb^sw^) applied to n = 12,282 transcripts) (Figure 5B), with a substantial increase of expression levels of *HBB, HBG1*, and *HBG2* (Figure 5C). Subsequently, we performed H3K9me3 ChIP-seq and found a notable reduction of H3K9me3 deposition on *HBB* and *HBG2* upon *WAVE2* deletion (Figure 5D). We then analyzed genome-wide H3K9me3 modification and observed a significant decrease of H3K9me3 deposition upon *WAVE2* deletion (Figure 5E), with consistent increase of CTCF and Rad21 enrichments (Figure 5F-I and S5C,D). Finally, we performed ATAC-seq experiments and found a significant increase of chromatin accessibility at *HBB, HBG1*, and *HBG2* (Figure 5J). Together, these data suggest that WAVE2 regulates *β-globin* gene expression via an H3K9me3-dependent mechanism.

**FIGURE 5.**
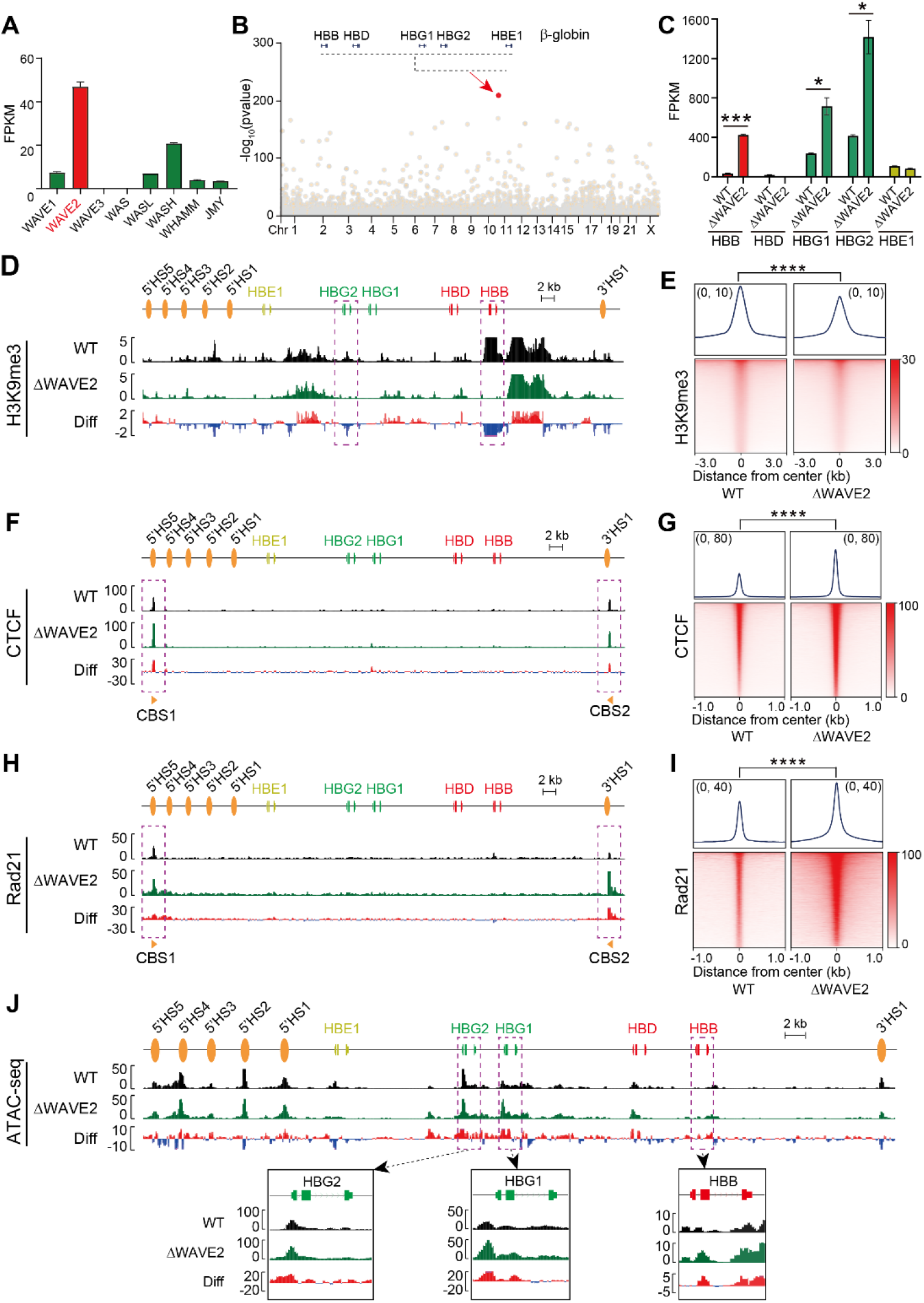
WAVE2 regulates *β-globin* gene expression. (A) Expression levels of the members of WASP family in K562 cells (n = 2). (B) Manhattan plots displaying localized enrichments (using 50kb sliding window) of differentially expressed genes induced by *WAVE2* knockout in K562 cells, with *β-globin* as a top-ranking locus. (C) RNA-seq data showing increased expression levels of *HBB, HBG1*, and *HBG2* upon *WAVE2* deletion in K562 cells (n = 2). (D) ChIP-seq profiles showing the loss of H3K9me3 modification on *HBB* and *HBG2* upon *WAVE2* deletion in K562 cells. (E) Aggregated peak analyses showing a significant decrease of H3K9me3 modification upon *WAVE2* deletion in K562 cells (n = 66,448). (F) ChIP-seq profiles showing increased CTCF enrichments at the two CBS elements flanking the *β-globin* gene cluster. (G) Aggregated peak analyses showing a significant increase of CTCF enrichments upon *WAVE2* deletion in K562 cells (n = 39,533). (H) ChIP-seq profiles showing increased Rad21 enrichments at the two CBS elements flanking the *β-globin* gene cluster. (I) Aggregated peak analyses displaying the significant increased Rad21 enrichments upon *WAVE2* deletion in K562 cells (n = 26,537). (J) ATAC-seq profiles showing increased DNA accessibility at *HBB, HBG1*, and *HBG2* in ΔWAVE2 cells compared to WT cells. Data are mean ± S.D, \**p* ≤ 0.05, \*\*\**p* ≤ 0.001, and \*\*\*\**p* ≤ 0.0001; unpaired two-tailed Student’s *t* test. FPKM, fragments per kilobase of exon per million mapped reads; Diff, difference (deletion vs WT).

## Discussion

Clustered protocadherins (cPcdhs) encoded by three tandem gene clusters—*Pcdh α, β*, and *γ*—are essential for neuronal identity and circuit assembly [25, 42, 43]. Each isoform is expressed from an individual promoter linked with a variable exon *cis*-spliced to a shared set of constant exons, generating a diverse repertoire of cell-surface proteins through stochastic yet patterned gene expression [25, 27, 44]. This unique variable and constant genomic organization is coupled to a regulatory mechanism involving long-range chromatin interactions: CTCF-binding sites within promoters and distal enhancers engage in cohesin-dependent loop extrusion to mediate selective promoter–enhancer spatial contacts, thereby governing promoter choice [19, 31-37]. The resulting combinatorial expression of *cPcdh* isoforms generates a molecular barcode that enables self-avoidance and self/non-self recognition among neurons, a process critical for dendritic arborization and the establishment of precise neural connectivity [38, 41].

The *cPcdh* locus is governed by higher-order chromatin mechanisms, including heterochromatin-mediated silencing and nuclear compartmentalization, highlighting its value as a model for understanding the interplay between genome architecture and stochastic gene regulation [24, 35, 46]. Upon release from heterochromatic silencing, individual *Pcdhα* promoters become accessible to CTCF binding, facilitating transcriptional activation through CTCF/cohesin-mediated promoter–enhancer looping. In addition, the transcriptionally inactive adult hemoglobin gene *HBB* localizes to the nuclear periphery, where it is thought to associate with the nuclear lamina in a heterochromatin state [62]. Moreover, nuclear actin polymerization has emerged as a key contributor to nuclear architecture, transcriptional regulation, and genome stability [13, 14, 63]. Within this framework, our findings identify WAVE2 as a regulator of *Pcdhα* expression through its role in maintaining heterochromatin. We show that WAVE2 regulates both stochastic and constitutive isoform expression of members of the *Pcdhα* cluster, linking actin regulatory machinery to higher-order chromatin states. In particular, ablation of *WAVE2* leads to expression switch from alternate members to c-type *Pcdhα* genes, concurrent with similar switch of chromatin accessibility at their promoters and the distal *HS5-1* enhancer (Figure 3F-L). This is consistent with cohesin-mediated nested loop extrusion in the opposite direction of convergent tandem CTCF sites within the *Pcdh* clusters [33, 35, 36]. Notably, *REST/NRSF* exerts a similar heterochromatin-dependent effect on this locus, suggesting that distinct pathways converge on a common chromatin-based Extending beyond regulatory *cPcdh*, mechanism. *WAVE2* also modulates perinuclear *β-globin* gene expression by influencing H3K9me3 depositions, suggesting a broader role in the regulation of clustered genes.

Together, these findings position *WAVE2* as a critical factor coupling actin dynamics to heterochromatin maintenance and genome regulation. Given prior evidence that Pcdh*α* and Pyk2 signaling act through the WAVE complex to control neuronal migration and cytoskeletal dynamics [12, 64, 65], it is tempting to speculate that feedback mechanisms may link cell-surface signaling to chromatin regulation. In this model, WAVE2 could serve as a molecular bridge between cytoskeletal signaling and 3D genome organization, providing a conceptual framework for how extracellular cues are integrated into nuclear gene regulatory programs. However, the precise molecular mechanisms by which *WAVE2* regulates heterochromatin organization and whether this function involves direct nuclear actin polymerization remain to be determined. Elucidating how *WAVE2* interfaces with chromatin and nuclear architecture will provide important insight into how cytoskeletal signaling pathways are integrated into genome regulation.

## Acknowledgments

We are grateful for advice on bioinformatics from Dr. Y. Zhang and discussion from all members of our laboratory. This study was supported by grants to Q.W. from the National Natural Science Foundation of China (32330016) and the National Key R&D Program of China (2022YFC3400200), and the grant to Y.T. from the National Natural Science Foundation of China (32200420).

## Author contributions

Q.W. conceived the research. L.W. and Y.T. performed experiments. L.W., Y.T. and H.H. analyzed data. L.W., Y.T., H.H. and Q.W. wrote the manuscript. All authors have read and approved the final manuscript.

---

**FIGURE S1.**
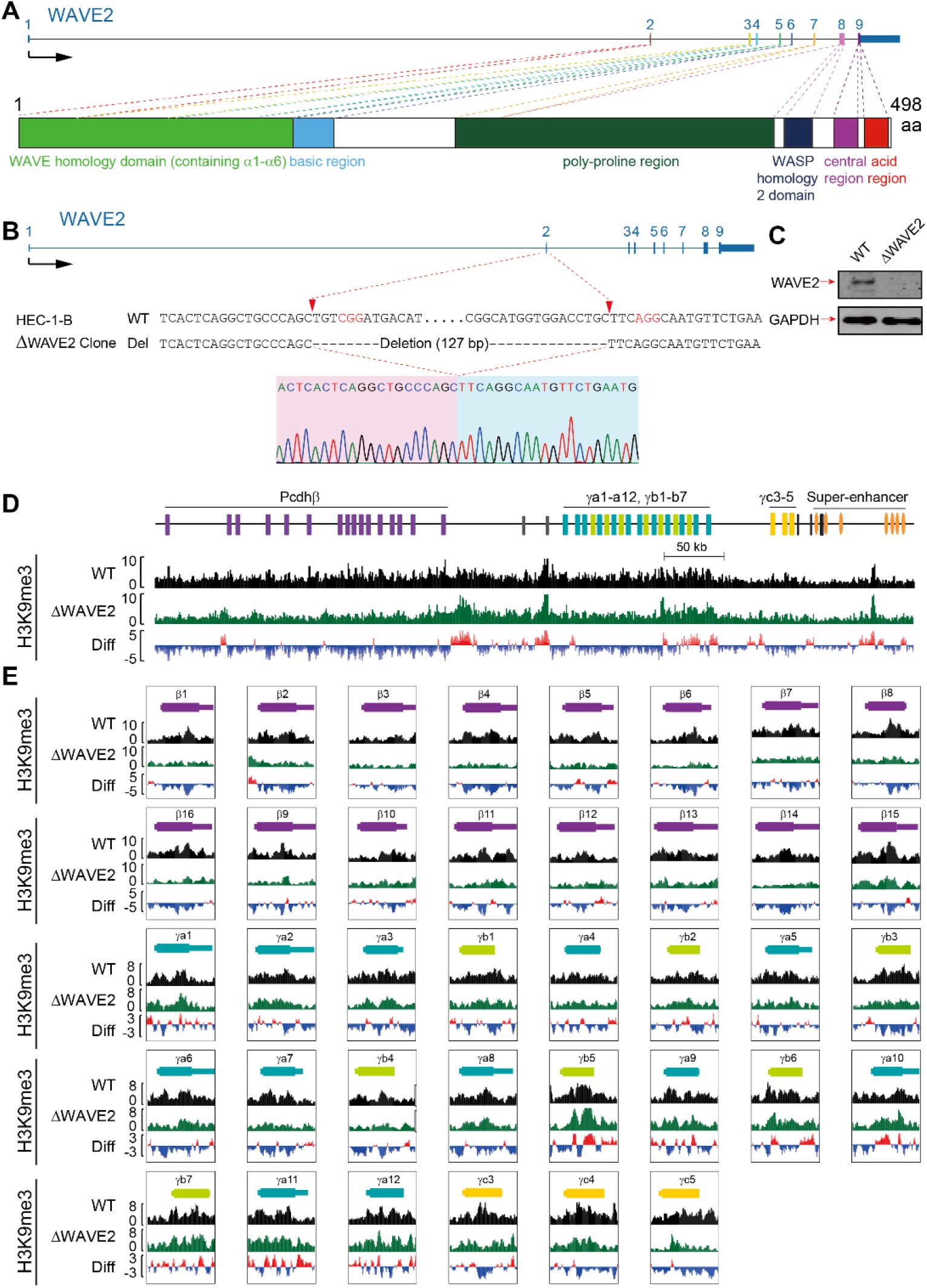
*WAVE2* deletion decreases H3K9me3 modification at *Pcdh β* and *γ* gene clusters in HEC-1-B cells. (A) Schematic of *WAVE2* genome organization (up) and the encoded protein domains (down). (B) Sanger sequencing confirmation of the *WAVE2*-knockout clone (ΔWAVE2) in HEC-1-B cells. (C) Western blot for the *WAVE2*-knockout HEC-1-B clone. (D and E) ChIP-seq profiles showing decreased H3K9me3 deposition on members of the *Pcdh β* and *γ* gene clusters in ΔWAVE2 and WT HEC-1-B cells. Diff, difference (deletion vs WT).

**FIGURE S2.**
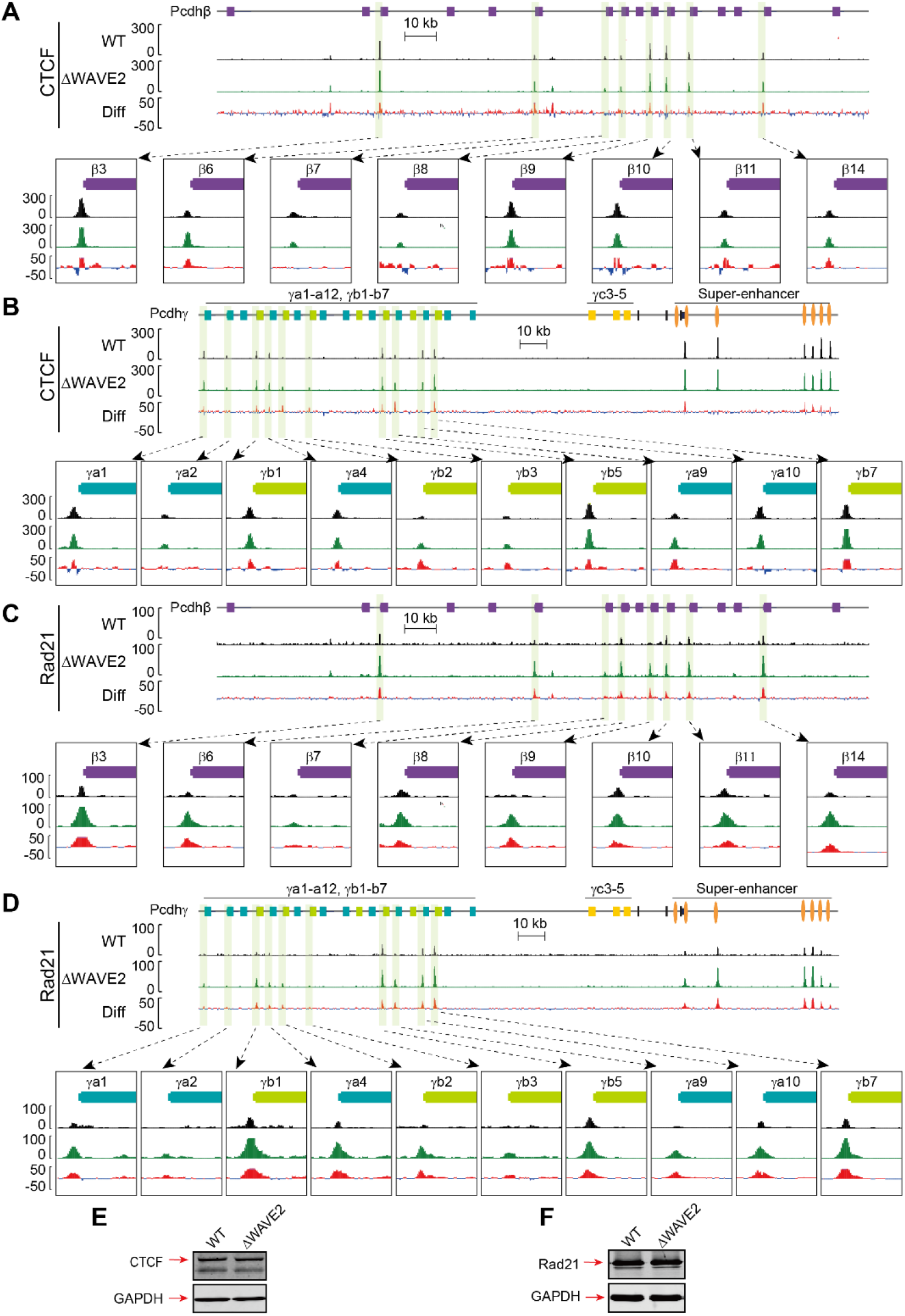
Loss of WAVE2 alters CTCF/cohesin binding profiles at the *Pcdh β* and *γ* gene clusters. (A and B) ChIP-seq profiles of CTCF at the *Pcdhβ* (A) and *Pcdhγ* (B) gene clusters in WT and ΔWAVE2 HEC-1-B cells. (C and D) ChIP-seq profiles of Rad21 at the *Pcdhβ* (C) and *Pcdhγ* (D) gene clusters in WT and ΔWAVE2 HEC-1-B cells. (E and F) Western blot showing expression levels of CTCF (E) and Rad21 (F) proteins in WT and ΔWAVE2 HEC-1-B cells. Diff, difference (deletion vs WT).

**FIGURE S3.**
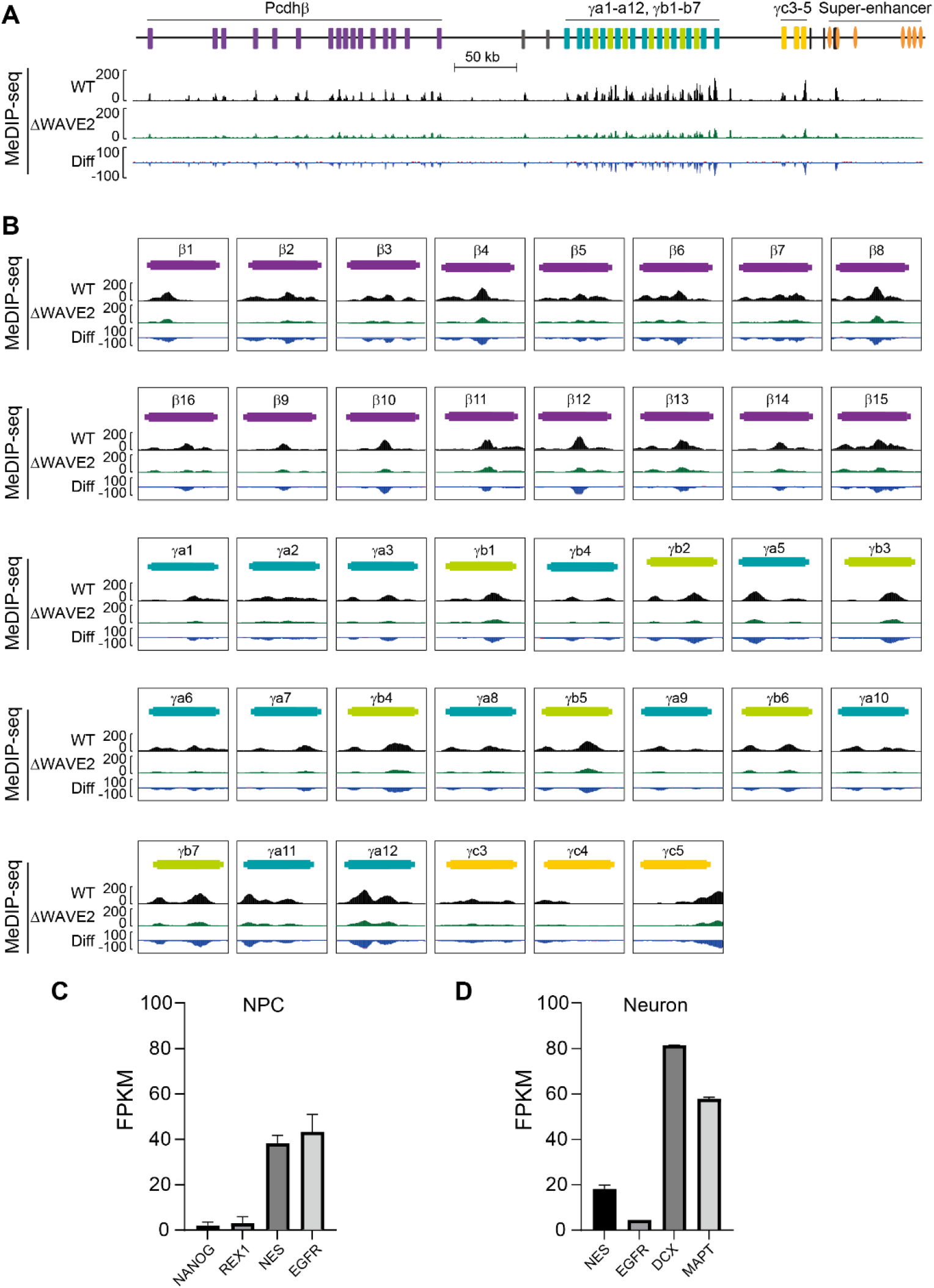
WAVE2 regulates DNA methylation at the *Pcdh β* and *γ* clusters. (A and B) MeDIP-seq showing decreased DNA methylation at members of the *Pcdh β* and *γ* gene clusters in ΔWAVE2 cells compared to WT cells. (C and D) Expression levels of *Nanog, Rex1, Nes*, and *Egfr* in NPCs (C), and *Nes, Egfr, Dcx, Mapt* in neurons (D). *Nanog* and *Rex1* are markers for ESC; *Nes* and *Egfr* are markers for NPC; *Dcx* and *Mapt* are markers for neuron. Diff, difference (deletion vs WT); FPKM, fragments per kilobase of exon per million mapped reads.

**FIGURE S4.**
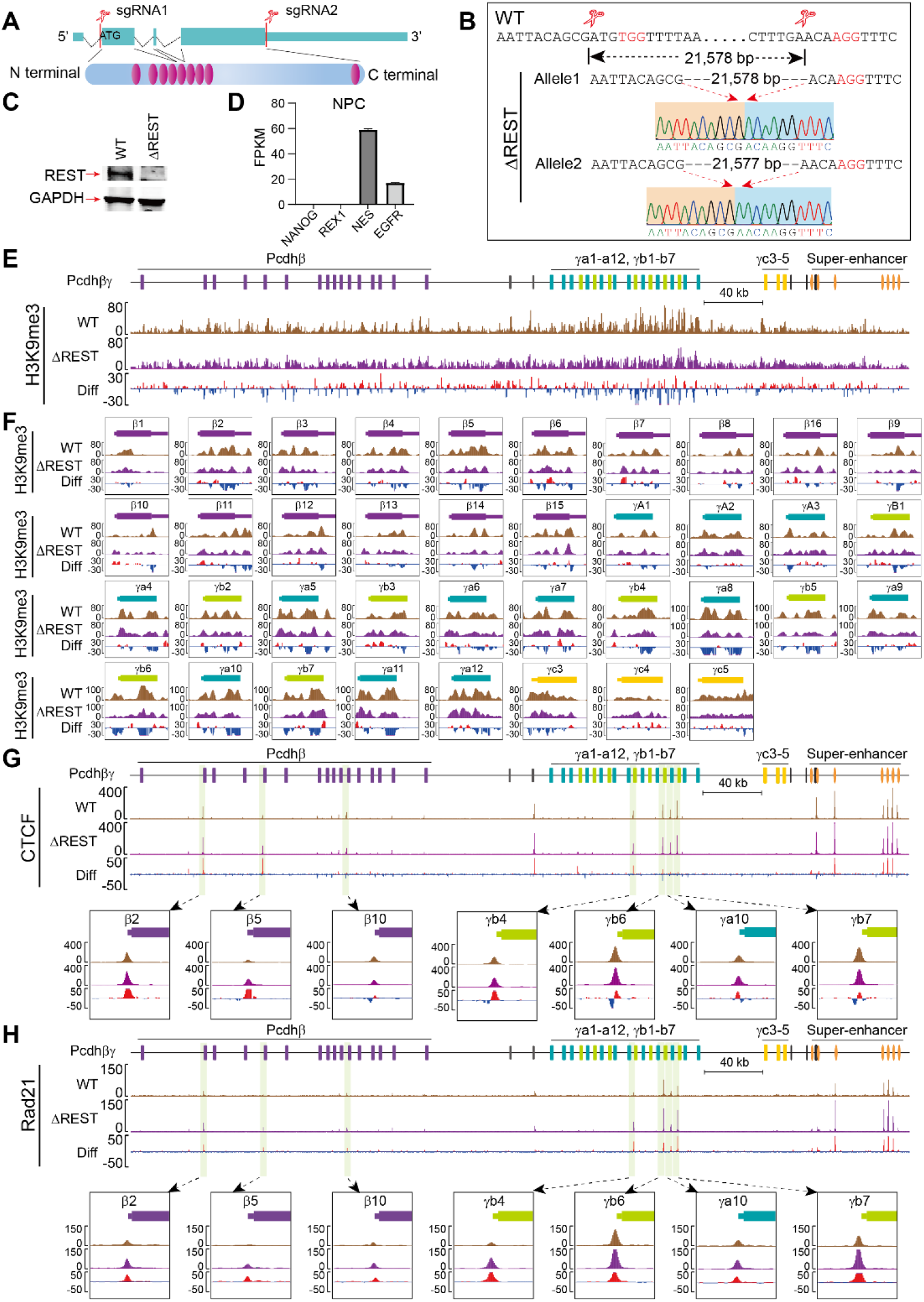
Deletion of *REST* affects the chromatin organization of the *Pcdh β* and *γ* gene clusters. (A) Schematic of the *REST* gene deletion showing genomic organization (up) and the encoded protein domains (down). The purple ovals represent the zinc-finger domains. (B) Sanger sequencing confirmation of the REST-knockout clone in human ESCs. ΔREST, the *REST* knockout clone, WT, the control clone. (C) Western blot of NPCs differentiated from the REST-knockout ESC clones. (D) Expression levels of *NANOG, REX1, NES* and *EGFR* in human NPCs. *NANOG* and *REX1* are markers for ESC; *NES* and *EGFR* are markers for NPC. (E and F) ChIP-seq profiles of H3K9me3 at *Pcdh β* and *γ* gene clusters in WT and ΔREST NPCs. (G) ChIP-seq profiles showing increased CTCF enrichments at *Pcdh β* and *γ* gene clusters in ΔREST compared to WT NPCs. (H) ChIP-seq profiles showing increased Rad21 enrichments at *Pcdh β* and *γ* gene clusters in ΔREST compared to WT NPCs. FPKM, fragments per kilobase of exon per million mapped reads; Diff, difference (deletion vs WT).

**FIGURE S5.**
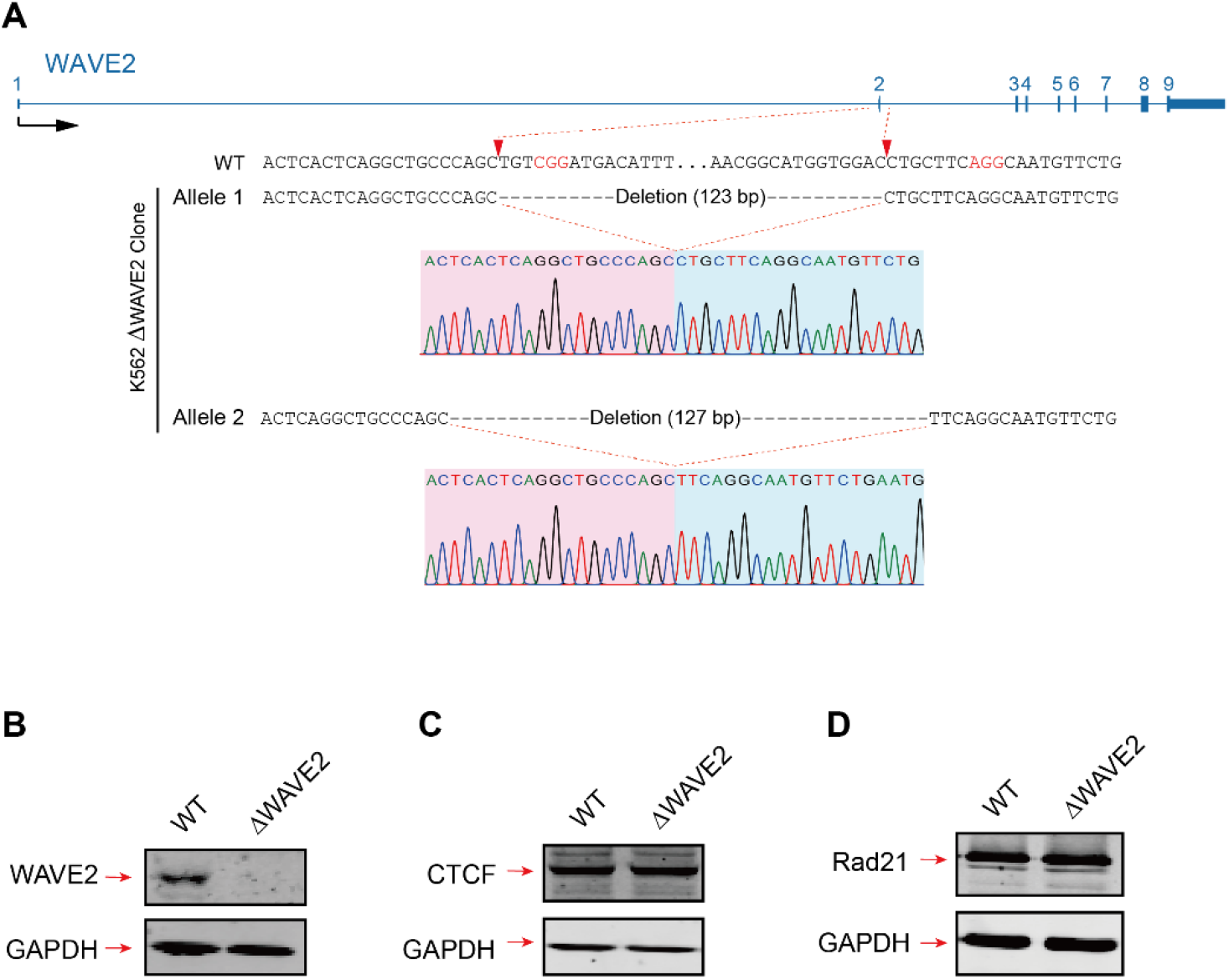
Knockout of *WAVE2* in K562 cells. (A) Sanger sequencing confirmation of the WAVE2-knockout clone in K562 cells. (B) Western blot for WAVE2 in K562 cells. (C and D) Western blots of the protein levels of CTCF (C) and Rad21 (D) in WT and ΔWAVE2 K562 cells.

**Supplementary Table 1.**
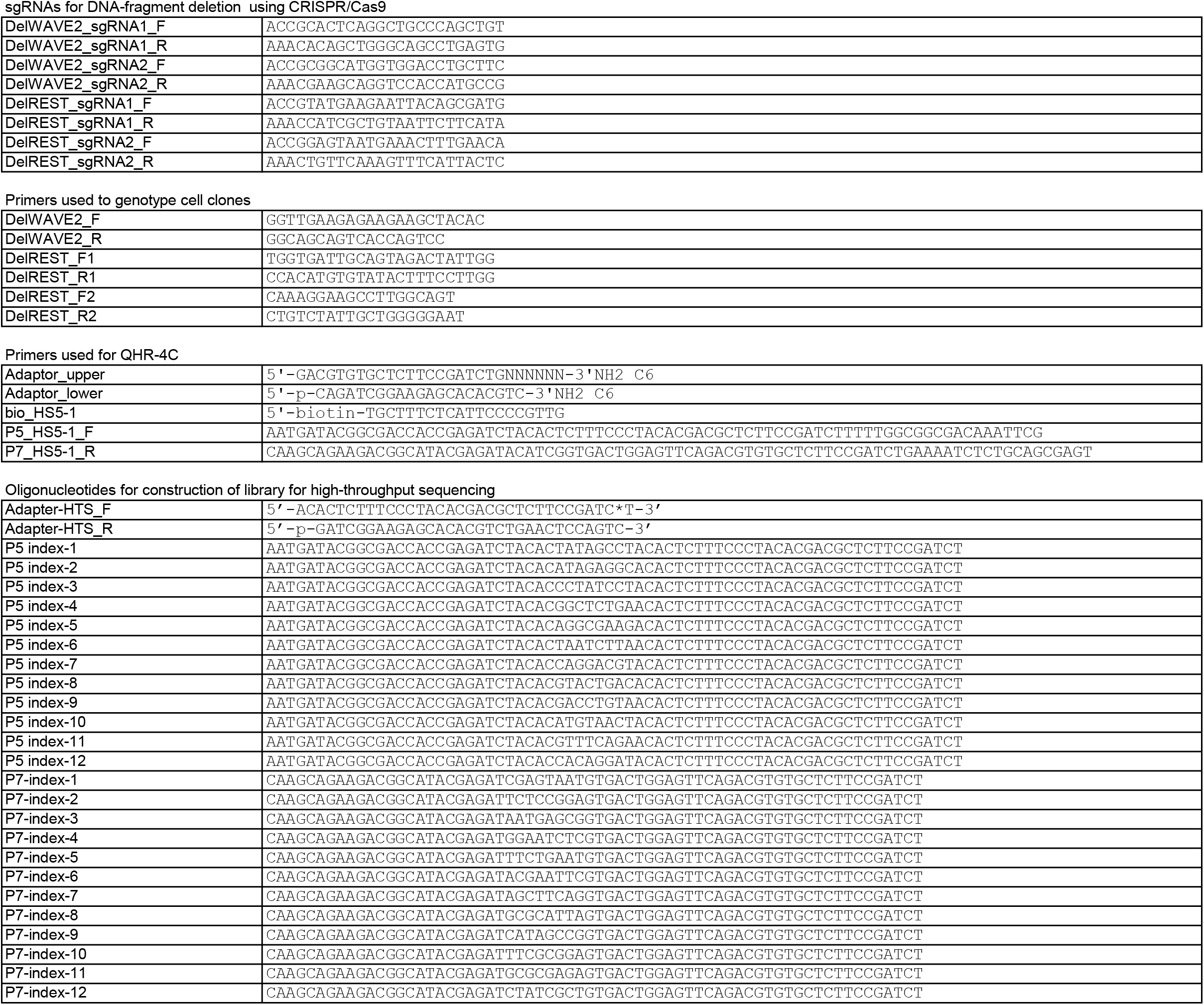

